# Taxonomic identification from metagenomic and metabarcoding data using any genetic marker

**DOI:** 10.1101/253377

**Authors:** Johan Bengtsson-Palme, Rodney T. Richardson, Marco Meola, Christian Wurzbacher, Émilie D. Tremblay, Kaisa Thorell, Kärt Kanger, K. Martin Eriksson, Guillaume J. Bilodeau, Reed M. Johnson, Martin Hartmann, R. Henrik Nilsson

## Abstract

Correct taxonomic identification of DNA sequences is central to studies of biodiversity using both shotgun metagenomic and metabarcoding approaches. However, there is no genetic marker that gives sufficient performance across all the biological kingdoms, hampering studies of taxonomic diversity in many groups of organisms. We here present a major update to Metaxa2 (http://microbiology.se/software/metaxa2/) that enables the use of any genetic marker for taxonomic classification of metagenome and amplicon sequence data.

---

Sequencing of DNA has revolutionized taxonomy, providing unprecedented resolution for species identification and definition^1, 2^. Similarly, the advent of large-scale sequencing techniques has opened entirely new windows on ecology, both for microbes and multicellular species^3^. In particular, high-throughput assignment of species and genus designations based on mixed samples of organisms or environmental substrates, so called DNA metabarcoding^4^, has made it possible to perform fine-tuned investigations of taxonomic diversity and to understand ecological interactions in different types of environments. However, an important bottleneck in such analyses is the size and quality of the reference sequence data to which the newly generated sequence reads are compared^5, 6^. Furthermore, while the ribosomal small-subunit (16S/18S/SSU) is a popular marker choice, no single genetic marker seems to be sufficient for covering all taxonomic groups with satisfactory accuracy for species or even genus assignments^7-9^. This has led to the establishment of a wide range of other genetic markers for DNA barcoding and metabarcoding in different organisms, such as *rbcL, matK, trnL*, and *trnH* for plants^10^, the ITS region for fungi^11^, and the COI gene for animals^12^. This broad diversity of DNA barcodes challenges sequence classification tools, which usually have been developed with the rRNA genes in mind^13-15^. Although some of these software tools can be re-trained on other reference dataseis, or have their reference databases exchanged for datasets representing other genes, they still make assumptions with regards to the reference data – such as global alignability – that often negatively affect performance, or prevent software operation altogether. In addition, increasing stringency with regards to correct taxonomic assignment often comes at the cost of lower proportions of classified sequences^16^. This tendency has been shown for some taxonomic classifiers also when operating on the rRNA genes^17^. The classification tool that appear least prone to show such a relationship is Metaxa2, which is based on a combination of hidden Markov models (HMMs) and sequence alignments^17^. Metaxa2 examines arbitrary DNA sequence datasets, such as genomes, metagenomes, or amplicons, and extracts the SSU and/or LSU rRNA genes; classifies the sequences to taxonomic origin; and optionally computes a range of diversity estimates for the studied community. However, Metaxa2 has so far been strictly limited to operation on the rRNA genes, preventing its use for other DNA barcodes. Yet, the capability of Metaxa2 to achieve high precision for its classifications while maintaining relatively high sensitivity would be highly desirable also for other genetic markers, particularly as these genes often are under-sampled in terms of species coverage^16^. Against this backdrop, the aim of this study was to adapt the Metaxa2 software for any additional DNA barcode. To this end, the paper presents an update to Metaxa2 itself, allowing the use of custom databases. We also introduce the Metaxa2 Database Builder – a software tool that allows users to create customized databases from DNA sequences and their associated taxonomic affiliations – and a repository for additional reference sets to meet the needs of the user.

The Metaxa2 Database builder has three different operating modes. The divergent mode is adapted to deal with barcoding regions for which fairly large sequence variability occurs among the target taxa, such as the eukaryotic ITS region^18^, the *trnH* gene used in plant barcoding^16^ and the COI gene used, e.g., for insects^12^. The conserved mode, on the other hand, is suitable for barcoding regions that are highly conserved among the target taxa, such as the SSU rRNA genes^19^ and the bacterial *rpoB* gene^20^. In addition, this mode is advisable for certain barcoding genes used in narrower taxonomic groups, such as Oomycota. In the conserved mode, the software extracts the barcoding regions from every input sequence and aligns them in order to determine the level of conservation across every position in the alignment. The most conserved regions are then extracted from the alignment and used to build HMMs that can be used to extract the barcoding region from metagenomic data. In the divergent mode, the database builder instead clusters the input sequences, aligns every individual cluster and builds one HMM for each cluster. The third mode – the hybrid mode – combines the features and advantages of the two others, but also their drawbacks. It should therefore only be used when none of those produces satisfactory results.

A key component for the high accuracy of Metaxa2 is the hand-curated classification database^17^. In the database builder, we have tried to emulate this curation by automating as much of our procedure as possible. There are three ways in which the software attempts to improve the taxonomic information. First, it can remove uninformative sequences from unknown specimens or mixed environmental samples. Second, it can make an effort to standardize the input taxonomy into seven levels. Finally, it can filter out entries without taxonomic affiliation at, for example, the genus or species level.

We evaluated the Metaxa2 Database Builder on 11 different barcoding regions, targeting a variety of uses (Supplementary Table 1). We first assessed the software performance on full-length sequences using the self-evaluation function, measured in terms of sensitivity, specificity, and error per assignment rate (Supplementary Fig. 1, see methods for details). In general, we found that at least one of the methods produced more than 80% correct assignments at the family level for half of the markers (Fig. 1a). However, three of the genetic markers – *rpb1, rpb2* and *cpn60* – consistently showed lower performance across all groups, even at the order level. When we multiplied the proportion of correct assignments with the total proportion of sequences assigned, it was clear that the divergent mode consistently was the best performing setting by this measure (Fig. 1b), mostly because the divergent mode always included the largest proportion of the input sequences in the final database (Supplementary Fig. 2). However, since the divergent mode includes essentially all input sequences in the classification database, it necessitates more careful manual curation of the dataset used for database creation. Therefore, if the data at hand is of uncertain quality, it may still be more adequate to use the conserved mode.

**Fig. 1.**
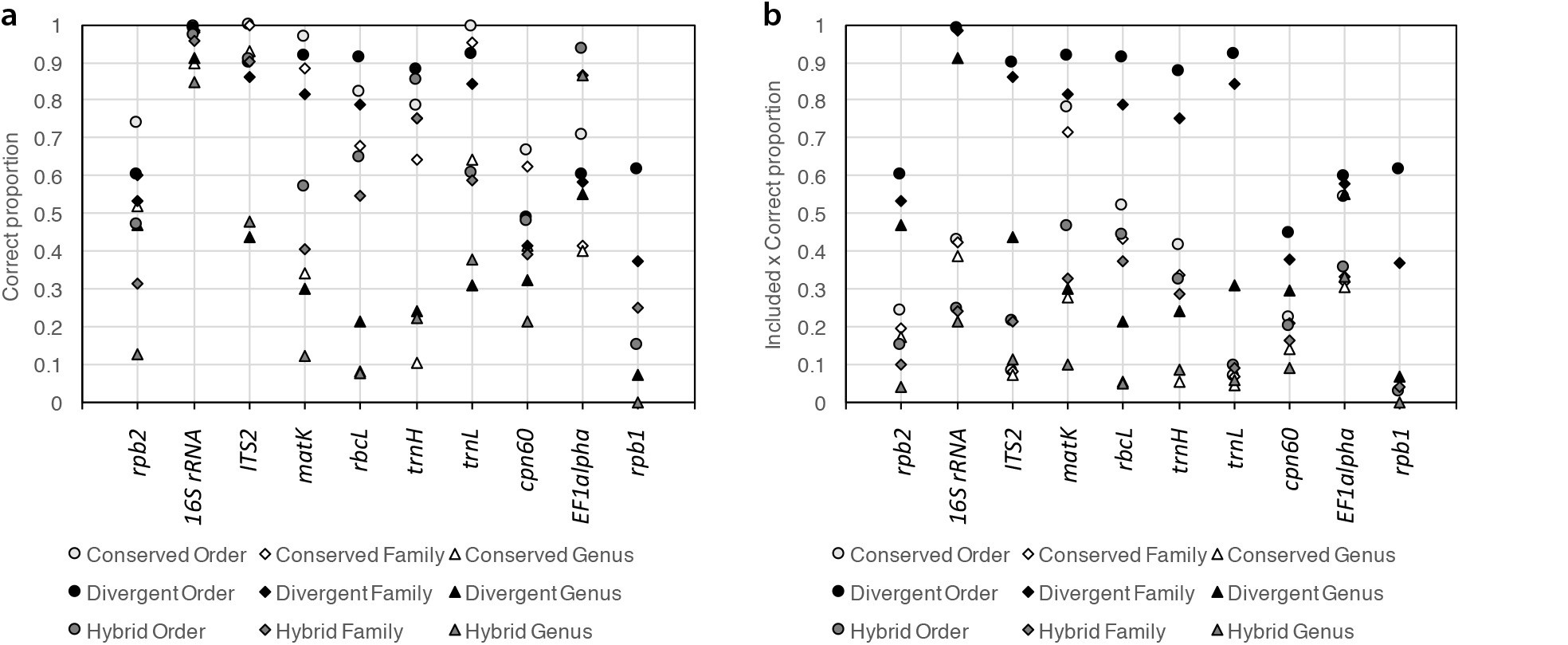
Self-evaluated performance of the Metaxa2 Database Builder in different operating modes (conserved, divergent and hybrid) on ten different DNA barcoding regions. A) Proportion of assigned sequences classified to the correct order (circles), family (diamonds) and genera (triangles). B) Proportion of correctly assigned sequences multiplied with the proportion of sequences included in the final classification databases (see Supplementary Fig. 2). The ATP9-NAD9 genetic marker is not shown, because it only had relevant taxonomic differences at the species level.

As an additional performance assessment, we followed the procedure from the original Metaxa2 evaluation^17^ and generated fragments of 150 nucleotides from each barcoding region to estimate the performance on shotgun metagenomic data. Here, we found that for most regions, the divergent mode generated the highest proportion of correct classifications (Fig. 2a). For EF1alpha, the hybrid mode performed better, for *matK* the operating modes were essentially tied, and for ATP9-NAD9 the conserved and hybrid modes performed the best. However, the divergent mode also produced higher numbers of misclassifications than the conserved mode did for ITS2, *matK* and *rbcL*, although the hybrid mode showed the largest numbers of incorrect assignments overall (Fig. 2b). Generally, the divergent mode showed the lowest levels of unclassified input sequences and over-predictions (Fig. 2c, 2d). Still, there are obvious differences in performance between different genetic markers. Particularly, it seems to be difficult to build appropriate models for the *rpb* genes and *cpn60*, at least based on the sequence data we used. Depending on what the user values the highest (comprehensiveness, stringency, precision etc.), different settings would be desirable, and several combinations of modes and filtering options should be evaluated against each other to find the optimal settings for each genetic marker and reference dataset. We furthermore compared the evaluation of the fragments to the internal software evaluation for each dataset (Supplementary Fig. 3). We found an essentially linear relationship between the proportion of sequences included in the database times the proportion of correct sequences in the internal evaluation and the proportion of correctly assigned sequence fragments (Supplementary Fig. 3e), and thus this may provide a robust measure of overall database performance.

**Fig. 2.**
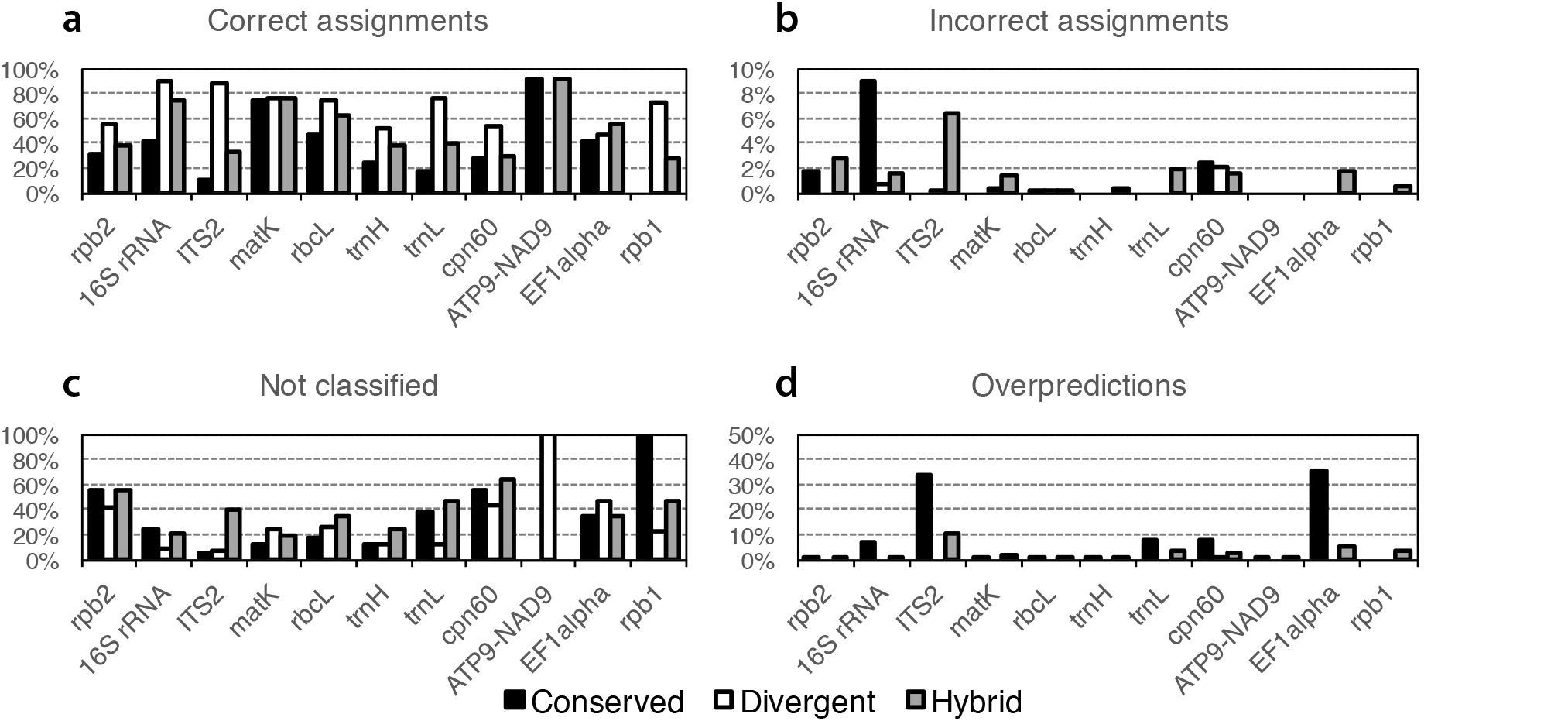
Family level Metaxa2 performance on randomly generated 150 bp fragments originating from the sequence datasets used to build the respective databases in the three different modes (Conserved, Divergent and Hybrid). A) Proportions of fragments assigned to the correct taxonomic family. B) Proportions of fragments assigned to an incorrect family. C) Proportions of fragments not assigned, or not recognized as belonging to the investigated barcoding region, at the family level. D) Family-level overpredictions, i.e. the proportions of sequence fragments belonging to a family not present in the final database, which were still assigned to a (different) family by Metaxa2. Note that the ATP9-NAD9 dataset is only used for species identification and thus this marker would be expected to show perfect performance on the family level. Note also that the Y-axis scales are different for B and for D compared to A and B.

We also compared the classification performance of the native Metaxa2 database to those resulting from automated construction based on SILVA. We built databases from release 111 (which was used as a starting point for the native Metaxa2 database) and release 128 in the conserved mode, with two versions for each release; one in which no filtering was applied and one in which we applied the filtering designed to mimic the manual curation process. We then classified simulated SSU fragments using Metaxa2, replacing the native database with the newly built ones. Overall, the results were surprisingly similar (Supplementary Fig. 4), contrary to what was previously shown when the native database was replaced with the GreenGenes database^17^. Interestingly, there were also rather small differences between the non-filtered and the automatically filtered databases, although applying filtering increased the number of classified sequence fragments with full taxonomic annotation and lowered the proportion of incorrect assignments, particularly at short fragment lengths. This indicates that the automated approach to database building is feasible, at least when the underlying sequence and taxonomy data are of high quality.

Evaluations of which taxonomic classification tools show the most consistent performance in terms of sensitivity and specificity are still largely incomplete^21^, particularly for non-standard barcoding regions, but commonly used software for taxonomic assignment, such as the RDP Naïve Bayesian Classifier^13^ and Rtax^14^, have all been shown to perform subpar or inconsistently in different settings^15-17^. We believe that the lack of comprehensive evaluation does not excuse the use of methods that produce incorrect or irrelevant results. With decreasing cost of DNA sequencing and increasing use of shotgun metagenomics for studies of biological communities, these updates to the Metaxa2 software – vastly extending its capabilities to virtually any high-quality DNA barcode in use – will enable a leap forward for molecular ecologists and others in need of precise taxonomic assignment among groups of taxa that are not feasibly targeted by traditional barcoding markers.

## Methods

### Software implementation

The Metaxa2 Database Builder (metaxa2_dbb) is a command-line, open source, Unix/Linux tool implemented in Perl. The software requires, on top of Perl, the Metaxa2^17^, HMMER3^22^, NCBI BLAST^23^, and MAFFT^24^ software to be installed. In addition, USEARCH^25^ or VSEARCH^26^ is highly recommended for full functionality. In short, the metaxa2_dbb tool creates the hidden Markov models (HMMs) and BLAST reference databases required to build a custom Metaxa2 classification database. Importantly, metaxa2_dbb can be run in three different operating modes, depending on how similar the sequences in the reference database are to each other.

In the conserved mode, used when sequences have regions of relatively high sequence similarity, the software first identifies a suitable main reference sequence, either by user selection or by clustering the sequences at 80% identity using USEARCH, and then selecting the representative sequence of the largest cluster. Next, it uses the (5’) start and (3’) end of the main reference sequence to define which of the other sequences in the input dataset should be considered full-length, and extracts those regions using Metaxa2. Thereafter, the identified full-length sequences are aligned using MAFFT, and the regions outside of the start and end of the main reference sequence are trimmed away before re-aligning the trimmed sequences again. This alignment is then used to determine the degree of sequence conservation across the alignment, to identify the regions of high and low conservation. The conserved regions of the alignment are extracted and aligned individually using MAFFT. Those alignments are used to build separate HMMs for each conserved region with hmmbuild of the HMMER package. The full-length input sequences matching at least half of those HMMs are then used to build the BLAST database used for classification, and their sequence IDs are edited to be compatible with the Metaxa2 database structure.

In the divergent mode, the input sequences are first clustered into groups with at least 20% sequence identity using USEARCH. Each such cluster is then aligned separately using MAFFT. The alignments are subsequently split at the mid position (including gaps), and each pair of alignments is used to build two separate HMMs using hmmbuild. The input sequences matching at least one of those HMMs are then used to build the BLAST database for classification, and their sequence IDs are edited as above. The hybrid mode is a combination of the conserved and divergent modes, in which the database builder will cluster the input sequences at 20% identity using USEARCH, and then proceed with same approach as in the conserved mode on each resulting cluster separately.

From this point, the analysis proceeds identically for the three modes. The software reads taxonomy data in any of the following formats: ASN.1, NCBI XML, and INSD XML formats, as provided by GenBank^27^; FASTA format with taxonomy data as part of the sequence headers, as provided by the SILVA^28^ and Greengenes^29^ databases; and the Metaxa2 tabulated taxonomy format. Optionally, the taxonomy data can be filtered to exclude sequences from uncultured or unknown organisms or with low-resolution taxonomic annotation information. The sequence data and taxonomic information are subsequently crosschecked such that entries are only retained if both sequence and taxonomy data are present. The remaining sequences are then compiled into a BLAST database using formatdb or makeblastdb of the BLAST/BLAST+ packages. Thereafter, unless pre-determined sequence identity cutoffs are provided by the user, suitable identity thresholds for taxonomic assignments at different classification levels are automatically determined. This is done by aligning the sequences in the BLAST database using MAFFT and then calculating the pairwise percent identity within and between taxonomic groups (intra- and inter-specific sequence identity). The identity cutoff for each taxonomic level is then set to be below the lowest intra-specific pairwise identity and, if possible, above the highest inter-specific pairwise identity. The cutoff can never be set to be above 99% identity for any taxonomic level.

Finally, the metaxa2_dbb software can perform an optional database evaluation step, which is further described below. A more thorough description of the database construction process can be found in the software manual (Supplementary Item 1). It should also be noted that to make the Metaxa2 classifier more reliable across a variety of barcoding regions, we have modified the algorithm for assigning reliability scores (see the manual for details; Supplementary Item 1). These modifications in general have very little effect on SSU and LSU classifications, but can nevertheless result in slight differences when the same dataset is classified using this version of Metaxa2 and versions prior to 2.2.

### Automatic correction of taxonomic data

If the user chooses, metaxa2_dbb can attempt to adjust the supplied taxonomy data in order to better match the taxonomic levels to those proposed by the Metaxa2 software (domain, phylum/kingdom, class, order, family, genus, species, and strain/subspecies). The phylum level is sorted out first, by checking which input taxonomic level that corresponds to a list of recognized phyla/kingdoms. This is followed by searching for a taxonomic level below the phylum level with an annotation ending with “-ales” to define the order level (unless the entry seems to be of metazoan origin). Then, the class level is defined as the level above the order level, and the family level is defined as the first level below the order level and with an annotation ending with “-ceae” (or “-idae” for metazoans). The species level is then identified by finding a taxonomic annotation similar to a Latin binomial using regular expressions. The genus level is finally defined as the level containing the genus part of the Latin binomial. This procedure can correct the vast majority of inconsistent taxonomic annotation data, although manual curation of the output data is highly recommended to catch exceptional cases.

### Use cases and software evaluation

We evaluated the metaxa2_dbb software by providing 12 different use cases involving 11 different DNA barcodes used in different scenarios (Supplementary Table 1). Notably, the datasets used to evaluate the software were not collected for the specific purpose of this evaluation, but were rather typical representatives of reference datasets used in previous or ongoing studies, thereby representing realistically relevant use cases for the Metaxa2 Database Builder very well. For the ITS2, *matK, rbcL, trnL* and *trnH* genetic markers, references were obtained from Richardson et al. (2017)^16^. Briefly, all NCBI nucleotide sequences for vascular plant available on 2016-03-04 were downloaded, filtered by length, and all sequences with more than two sequential uncalled nucleotides were removed. The datasets were then filtered to remove duplicates and sequences from plants not present in Ohio and surrounding states and provinces. Taxonomic information was obtained from NCBI taxonomy^30^. Sequences with undefined taxonomic information at any rank were removed. For *rpb1, rpb2* and *EFalpha*, references were obtained from the fungal six-gene phylogeny of James et al.^31^. Sequence data and taxonomic information were obtained from NCBI. For the 16S rRNA gene, sequences and taxonomic data for type-strains and cultured strains were downloaded from SILVA release 128^32^, and SATIVA^33^ was used to remove mislabeled strains. For *cpn60*, sequences were downloaded from the cpnDB^34^ as of 2016-10-21. The complete nucleotide sequences of group I chaperonins, i.e. *cpn60* (also known as *hsp60* or *groEL*), which is found in bacteria, some archaea, mitochondria and plastids, were used for building the database. Two datasets were downloaded, both the FASTA file of all group I sequences and a reduced file with only reference genome representatives. Taxonomic classifications were transferred from the SILVA annotation of release 111 and then manually curated. Finally, for ATP9-NAD9, we used a database assembled from curated sequences including 140 different *Phytophthora* species/hybrids (GenBank accession numbers JF771616.1 to JF772053.1 and JQ439009.1 to JQ439486.1, and Bilodeau and Robideau^35^; n.b. a total of 123 species are currently described; http://www.phytophthoradb.org).

When sequence and taxonomic data had been obtained for each of these genetic markers, we ran the metaxa2_dbb software on each data set using the conserved, divergent and hybrid modes. We also enabled the self-evaluation option, which performs a cross-validation of the database performance similar to that of Richardson et al.^16^. For the self-evaluation we used the default settings, which correspond to rebuilding the database ten times, each time using 90% of the input sequences to build the database (the training set) and then subsequently classifying the remaining 10% of input sequences (the testing set) using Metaxa2. The predicted taxonomic classifications were then compared against the taxonomic identity of each test sequence dervied from the source databases at every taxonomic level, generating measures for sensitivity (proportion of test sequences identified as matching the barcoding region), specificity (proportion of correctly classified sequences at the taxonomic level in question), and the error per classification ratio (proportion of incorrectly classified sequences per total classifications made).

In addition to the software self-evaluation, we also tested the classification performance of the different databases on sequence fragments derived from the sequences used to build the respective database. This evaluation followed the method used for the original Metaxa2 paper^17^, although we only generated fragments of a single length, viz. 150 nucleotides. The test sets were generated by randomly selecting a stretch of 150 nucleotides from every sequence in the input data for each barcoding region. We then used Metaxa2 version 2.2 to classify these simulated read data sets and calculated the performance for each barcoding region in terms of accuracy (proportion of correctly classified sequence fragments), misclassifications (incorrect assignments), sensitivity (proportion of non-detected sequence fragments), and over-prediction (incorrect assignment to a rank for which there is no reference belonging to the query taxa present in the database). Sequence fragments were regarded as correctly classified if their reported taxonomy corresponded to the known taxonomy of the input sequence that the fragment was derived from, at every taxonomic level as reported by Metaxa2. If any incorrect taxonomic affiliations were reported at any taxonomic level, the fragment was regarded as misclassified.

We finally compared the performance of the hand-curated Metaxa2 SSU rRNA database that is bundled with the software to SSU rRNA databases built by metaxa2_dbb from the sequences in SILVA release 111 and 128^28^. The native Metaxa2 database is based on SILVA release 111, which means that the comparison between the native database and release 111 is relevant to understand the differences between the manual and automatic database constructions. The difference to release 128, on the other hand, is rather a test of whether the accuracy changes with the addition of more reference sequences. The SILVA databases were created by downloading the FASTA file representing the reference SSU sequences with 99% non-redundancy (SSURef_Nr99) with taxonomy from SILVA. We then added the SSU sequences for the 12S rRNA used in the native Metaxa2 database from MitoZoa^17, 36^. From these, we used Metaxa2 version 2.1.2 (default settings) to divide the SSU sequences by taxonomic domain. The resulting files were used as input for metaxa2_dbb, which was run by retaining the HMM profiles from the native database, i.e. only rebuilding the classification database. In all cases, taxonomy correction was used, and cutoffs were manually set to “0,60,70,75,85,90,97”^17^. The full options were: “metaxa2_dbb -o SSU_SILVAXXX -g SSU -p metaxa2_db/SSU/HMMs/ -t SILVA_XXX_SSURef_Nr99_tax_silva.fasta -a archaea.fasta -b bacteria.fasta -c chloroplast.fasta -e eukaryota.fasta -m mitochondria.fasta -n mitozoa_SSU.fasta –correct_taxonomy T –cutoffs ‘0,60,70,75,85,90,97’ –cpu 16”. For each SILVA release, two databases were built, one with the command above, and one in which filtering of taxonomic information was applied, adding the “–filter_uncultured T –filter_level 6” options.

After these new SILVA-based classification databases had been constructed, we classified the simulated SSU read fragments with high-quality taxonomic information used in the original Metaxa2 evaluation, and ran this in the same way as in the original paper^17^. The results of the classifications were investigated manually to make sure that errors made by Metaxa2 were due to actual classification errors and not renaming of taxa, inconsistencies in taxonomy between database versions, synonymous names used for one taxon, or misspellings. As in the original Metaxa2 paper, a sequence fragment was regarded correctly classified if the reported taxonomy corresponded to the known taxonomy of the input sequence at every taxonomic level, as reported by Metaxa2. If the Metaxa2 classification was found to completely correspond to the known taxonomic affiliation at all investigated taxonomic levels, the sequence fragment was regarded as perfectly classified. If Metaxa2 reported any incorrect taxonomic affiliation at any taxonomic level the fragment was regarded as misclassified.

## Acknowledgements

The authors would like to thank Prof. Christer Erséus for input on the International Code of Zoological Nomenclature. JBP acknowledges financial support from the Swedish Research Council for Environment, Agricultural Sciences and Spatial Planning (FORMAS; grant 2016-768). RTR was supported by a Project Apis m. - Costco Honey Bee Biology Fellowship. KME acknowledges financial support from FORMAS (grant 2012-86). RHN acknowledges funding from FORMAS (grant 215-2011-498).

## Author contributions

JBP and RTR conceived and designed the study. JBP designed the software, with input from RTR, RMJ, MH and RHN. JBP implemented the software. JBP, KME and RHN updated the software manual. RTR, CW, EDT, MM, KT, GJB and RMJ provided and curated data for software evaluation. JBP, RTR, CW, MM, KT and KK evaluated software performance. JBP, KME, MH and RHN updated the classification databases. JBP drafted the manuscript, with help from RTR and RHN. All authors contributed to and approved the final manuscript.

## References

1. Woese, C. R., Kandler, O. & Wheelis, M. L. Towards a natural system of organisms: proposal for the domains Archaea, Bacteria, and Eucarya. Proc Natl Acad Sci USA 87, 4576–1579 (1990).

2. Hibbett, D. et al. Sequence-based classification and identification of Fungi. Mycologia (2016). doi:10.3852/16-130

3. Yoccoz, N. G. The future of environmental DNA in ecology. Mol Ecol 21, 2031–2038 (2012).

4. Taberlet, P., Coissac, E., Pompanon, F., Brochmann, C. & Willerslev, E. Towards next-generation biodiversity assessment using DNA metabarcoding. Mol Ecol 21, 2045–2050 (2012).

5. Bengtsson-Palme, J. et al. Strategies to improve usability and preserve accuracy in biological sequence databases. Proteomics 16, 2454–2460 (2016).

6. Nilsson, R. H. et al. Taxonomic reliability of DNA sequences in public sequence databases: a fungal perspective. PLoS ONE 1, e59 (2006).

7. Wang, X.-C. et al. ITS1: a DNA barcode better than ITS2 in eukaryotes? Mol Ecol Resour 15, 573–586 (2015).

8. Lindahl, B. D. et al. Fungal community analysis by high-throughput sequencing of amplified markers - a user’s guide. New Phytol (2013). doi:10.1111/nph.12243

9. Bruns, T. D. & Taylor, J. W. Comment on “Global assessment of arbuscular mycorrhizal fungus diversity reveals very low endemism”. Science 351, 826 (2016).

10. Richardson, R. T. et al. Rank-based characterization of pollen assemblages collected by honey bees using a multi-locus metabarcoding approach. Appl Plant Sci 3, (2015).

11. Schoch, C. L. et al. Nuclear ribosomal internal transcribed spacer (ITS) region as a universal DNA barcode marker for Fungi. Proc Natl Acad Sci USA 109, 6241–6246 (2012).

12. Hebert, P. D. N., Ratnasingham, S. & deWaard, J. R. Barcoding animal life: cytochrome c oxidase subunit 1 divergences among closely related species. Proc Biol Sci 270 Suppl 1, S96–9 (2003).

13. Wang, Q., Garrity, G. M., Tiedje, J. M. & Cole, J. R. Naive Bayesian classifier for rapid assignment of rRNA sequences into the new bacterial taxonomy. Appl Environ Microbiol 73, 5261–5267 (2007).

14. Soergel, D. A. W., Dey, N., Knight, R. & Brenner, S. E. Selection of primers for optimal taxonomic classification of environmental 16S rRNA gene sequences. ISME J 6, 1440–1444 (2012).

15. Edgar, R. SINTAX: a simple non-Bayesian taxonomy classifier for 16S and ITS sequences. bioRxiv 074161 (2016). doi:10.1101/074161

16. Richardson, R. T., Bengtsson-Palme, J. & Johnson, R. M. Evaluating and optimizing the performance of software commonly used for the taxonomic classification of DNA metabarcoding sequence data. Mol Ecol Resour 17, 760–769 (2017).

17. Bengtsson-Palme, J. et al. Metaxa2: Improved identification and taxonomic classification of small and large subunit rRNA in metagenomic data. Mol Ecol Resour 15, 1403–1414 (2015).

18. Nilsson, R. H. et al. Five simple guidelines for establishing basic authenticity and reliability of newly generated fungal ITS sequences. MycoKeys 4, 37–63 (2012).

19. Hartmann, M., Howes, C. G., Abarenkov, K., Mohn, W. W. & Nilsson, R. H. V-Xtractor: an open-source, high-throughput software tool to identify and extract hypervariable regions of small subunit (16S/18S) ribosomal RNA gene sequences. J Microbiol Methods 83, 250–253 (2010).

20. Dahllöf, I., Baillie, H. & Kjelleberg, S. rpoB-based microbial community analysis avoids limitations inherent in 16S rRNA gene intraspecies heterogeneity. Appl Environ Microbiol 66, 3376–3380 (2000).

21. Bengtsson-Palme, J. Strategies for Taxonomic and Functional Annotation of Metagenomes in Metagenomics: Perspectives, Methods, and Applications (ed. Nagarajan, M.) 55–79 (Academic Press, Elsevier, Oxford, 2018).

22. Eddy, S. HMMER. http://hmmer.janelia.org (2010).

23. Altschul, S. F. et al. Gapped BLAST and PSI-BLAST: a new generation of protein database search programs. Nucleic Acids Res 25, 3389–3402 (1997).

24. Katoh, K. & Standley, D. M. MAFFT multiple sequence alignment software version 7: improvements in performance and usability. Mol Biol Evol 30, 772–780 (2013).

25. Edgar, R. C. Search and clustering orders of magnitude faster than BLAST. Bioinformatics 26, 2460–2461 (2010).

26. Rognes, T., Flouri, T., Nichols, B., Quince, C. & Mahé, F. VSEARCH: a versatile open source tool for metagenomics. PeerJ 4, e2584 (2016).

27. Clark, K., Karsch-Mizrachi, I., Lipman, D. J., Ostell, J. & Sayers, E. W. GenBank. Nucleic Acids Res 44, D67–72 (2016).

28. Quast, C. et al. The SILVA ribosomal RNA gene database project: improved data processing and web-based tools. Nucleic Acids Res 41, D590–6 (2013).

29. McDonald, D. et al. An improved Greengenes taxonomy with explicit ranks for ecological and evolutionary analyses of bacteria and archaea. ISME J 6, 610–618 (2012).

30. Federhen, S. The NCBI Taxonomy database. Nucleic Acids Res 40, D136–43 (2012).

31. James, T. Y. et al. Reconstructing the early evolution of Fungi using a six-gene phylogeny. Nature 443, 818–822 (2006).

32. Yilmaz, P. et al. The SILVA and ‘All-species Living Tree Project (LTP)’ taxonomic frameworks. Nucleic Acids Res 42, D643–8 (2014).

33. Kozlov, A. M., Zhang, J., Yilmaz, P., Glöckner, F. O. & Stamatakis, A. Phylogeny-aware identification and correction of taxonomically mislabeled sequences. Nucleic Acids Res 44, 5022–5033 (2016).

34. Hill, J. E., Penny, S. L., Crowell, K. G., Goh, S. H. & Hemmingsen, S. M. cpnDB: a chaperonin sequence database. Genome Res 14, 1669–1675 (2004).

35. Bilodeau, G. J. & Robideau, G. P. Optimization of nucleic acid extraction from field and bulk samples for sensitive direct detection of plant pests. Phytopathology 104, S3.14 (2014).

36. D’Onorio de Meo, P. et al. MitoZoa 2.0: a database resource and search tools for comparative and evolutionary analyses of mitochondrial genomes in Metazoa. Nucleic Acids Res 40, D1168–72 (2012).

